# Diurnal rhythmicity modulates self-blaming behaviours and neural processing of autobiographical guilt memories in the trait-like and state-like manner

**DOI:** 10.64898/2026.09.22.753486

**Authors:** Michal Rafal Zareba, Ivan González-García, Marcos Ibáñez Montolio, Maya Visser

**Affiliations:** Neuropsychology and Functional Neuroimaging Group, Department of Basic and Clinical Psychology and Psychobiology, Jaume I University, 12071 Castellon de la Plana, Spain

**Keywords:** guilt, self-blame, chronotype, distinctness, diurnal rhythmicity, fMRI, anxiety

## Abstract

Increasing evidence implicates late chronotype and higher distinctness as two diurnal dimensions associated with a risk for affective dysregulation. Nonetheless, the underlying behavioural and neural mechanisms remain poorly understood. Previous works in depressive and anxious populations have demonstrated an important role of maladaptive self-blaming processes, with pivotal contributions from superior anterior temporal lobes (sATL), involved in social conceptual knowledge, and subgenual anterior cingulate cortex (sgACC), related to self-worth and social affiliative processing. In the current study, we first tested how diurnal rhythm characteristics were associated with self-blaming emotions and related coping mechanisms through hypothetical social scenarios where participants or their close friends acted towards one another against social norms. Secondly, to probe alterations in the neural correlates of self-blaming, the participants underwent functional neuroimaging during reliving of autobiographical guilt memories. On the behavioural level, late chronotypes demonstrated decreased self-distancing and contextually-maladaptive blaming patterns, i.e. increased blaming of the action recipients rather than the agents. This was paralleled by decreased self-blame-related activity in the sATL and sgACC. Distinctness was associated with altered task-dependent sATL connectivity with hippocampus, brainstem and cerebellum. Importantly, some of the neuroimaging findings reflected interactions between the diurnal rhythm characteristics and time-of-day. As such, the current work provides plausible correlates of adverse emotional outcomes in late chronotypes. Furthermore, it suggests that affective processing reflects an interplay between stable trait-like characteristics and dynamic state-like processes varying across the day, contributing to fine-grained understanding of how emotional cognition is embedded in the brain.

## 1. Introduction

Human functioning, including arousal and affect, exhibits diurnal variation driven by circadian and homeostatic sleep processes (Cajochen and Schmidt, 2025; Cox et al., 2024). Across development, wakefulness timing shifts from earlier hours in childhood to later chronotypes during adolescence and young adulthood (Randler et al., 2017), coinciding with a period of increased vulnerability to affective disorders (Taylor and Hasler, 2018). Longitudinal evidence suggests bidirectional links between circadian functioning and affective symptoms (Haraden et al., 2017), with the leading theories postulating that late chronotypes accumulate chronic sleep debt under socially-imposed schedules, impairing their emotion regulation abilities (Jamieson et al., 2021). Beyond chronotype, the subjective amplitude of diurnal rhythms (distinctness) has recently emerged as an independent factor. This measure of how much your state (e.g. alertness, mood, energy) varies across the day has been associated with negative emotionality and sleep problems (Carciofo and Song, 2019; Scislewska et al., 2025). However, it remains unknown whether the effects of chronotype and distinctness on sleep and mental health are co-dependent.

Neuroimaging literature on potential bases of these observations remains scarce. Late chronotypes show reduced medial prefrontal responses during reward anticipation (Hasler et al., 2013). Evening-types also display heightened amygdala responses to negative emotional faces, paralleled by weaker amygdala–dorsal anterior cingulate connectivity, suggesting impaired implicit emotion regulation (Horne and Norbury, 2018; Jamieson et al., 2021). Only a single work has investigated the role of distinctness, reporting positive associations with the activity of the ventral tegmental area and occipital pole during anticipation of punishment and negative feedback, respectively (Scislewska et al., 2025). Importantly, although these studies controlled for time-of-day (TOD) effects, they did not test for their potential interactions with diurnal rhythm characteristics. If chronotype determines the timing of individual affective rhythms, and distinctness describes their strength, these associations should in theory be reflected in the underlying neural networks.

To extend previous work from basic affective processing to more socially-demanding contexts, we focused on self-blaming emotions. Increased self-blaming and maladaptive ways of dealing with it, e.g. social distancing and self-criticism, are a common characteristic of depressive and anxious symptomatology (Cândea and Szentagotai-Tătar, 2018; Duan et al., 2023; Zareba et al., 2026). At the brain level, several neuroimaging works in patients and subclinical cohorts have suggested functional integration between superior anterior temporal lobes (sATL), involved in social conceptual knowledge (Thye et al., 2024), and subgenual anterior cingulate cortex (sgACC), linking self-worth and social affiliative processing (Zahn, 2025), as a promising biomarker of maladaptive self-blaming (Lythe et al., 2015; Zahn et al., 2019; Fennema et al., 2023; Zareba et al., 2024; Zareba et al., 2026).

Therefore, the present study had three aims. Firstly, we sought to verify whether distinctness moderated the associations of chronotype with sleep and mental health metrics, expecting higher distinctness to be linked to more adverse outcomes. Secondly, we used the Moral Sentiment and Action Tendencies (MSAT; Zahn et al., 2015; Duan et al., 2023) to characterise how diurnal rhythmicity was related to self-blaming emotions and associated action tendencies, hypothesising to observe similar patterns as those reported in depressed and subclinically anxious individuals (Duan et al., 2023; Zareba et al., 2026). Last but not least, we tested how diurnal factors were related to neural correlates of re-experiencing autobiographical guilt memories, i.e. an instance of self-blaming behaviours. The analysis considered both the average task effects and their habituation. We expected to observe main effects of chronotype and distinctness, akin to the previous studies (Hasler et al., 2013; Horne and Norbury, 2018; Scislewska et al., 2025), but also their interactions with TOD, reflecting trait-like and state-like processes, respectively.

## 2. Methods

The current work includes a battery of psychometric questionnaires, a behavioural task on self- and other-blaming emotions, and functional magnetic resonance imaging (fMRI) with a task on autobiographical guilt memory processing. The data was collected as a part of the project investigating self-blaming emotions in subclinically anxious individuals (Zareba et al., 2026). The original protocol adhered to the ethical standards of the Declaration of Helsinki and was approved by the local ethics committee (CEISH/07/2022). Participants provided informed consent prior to their participation and were remunerated at a rate of 10 euros per hour.

### 2.1. Participants and psychometric measures

165 healthy volunteers were recruited predominantly from the university community via local announcements and word of mouth. Given that all the participants were subsequently planned to undergo MRI, the typical inclusion criteria for MRI studies were applied: right-handedness (determined with Edinburgh Handedness Inventory; Oldfield, 1971), normal or corrected to normal vision, no history of neurological and psychiatric disorders, and no current psychotropic medication. Upon being invited to the laboratory, the participants completed a battery of psychometric questionnaires. These included measurements of trait-anxiety and concurrent depressive symptomatology, as determined with State-Trait Anxiety Inventory (STAI; Spielberger, 1989) and Beck Depression Inventory (BDI; Beck et al., 1961), respectively. Collected chronobiological and sleep variables included individual chronotype, distinctness and morning affect, assessed with the Morningness-Eveningness-Stability-Scale Improved (MESSi; Randler et al., 2016; Díaz-Morales and Randler, 2017), together with self-reported sleep need and sleep quality over the past month, the latter determined with Pittsburgh Sleep Quality Index (PSQI; Buysse et al., 1989). Based on the obtained information, two additional sleep-related measures were calculated: (1) reported sleep midpoint (PSQI), and (2) reported sleep debt, i.e. the difference between the sleep time (PSQI) and sleep need (a single item self-report).

During their visit to the laboratory, a subset of the participants (n = 140) additionally performed a computerised version of the Spanish adaptation of the MSAT (Zahn et al., 2015; Duan et al., 2023; Zareba et al., 2026). Furthermore, 73 individuals underwent fMRI during autobiographical guilt recollection task (Zareba et al., 2026; n = 73). The characteristics of the task-specific cohorts are presented in **Table 1**.

**Table 1.** Demographic and psychometric characteristics of the full cohort (n = 165), together with the subsamples who performed the Moral Sentiment and Action Tendencies task (MSAT; n = 140) and autobiographical guilt recollection task (n = 73; task fMRI). The upper index values indicate the number of participants for whom a given measure was missing.

| Variable | Mean ± standard deviation | Possible range |
| --- | --- | --- |
| Sex | Full cohort: 100 females, 65 males<br>MSAT: 87 females, 53 males<br>task fMRI: 46 females, 27 males | not applicable |
| Age | Full cohort: 22.55 ± 3.98<br>MSAT: 22.56 ± 4.07<br>task fMRI: 23.85 ± 4.74 | not applicable |
| Trait-anxiety (STAI) | Full cohort: 23.27 ± 12.24<br>MSAT: 23.31 ± 12.30<br>task fMRI: 23.44 ± 12.17 | 0-60 |
| Depressive symptoms (BDI) | Full cohort: 12.85 ± 10.30<br>MSAT: 13.35 ± 10.85<br>task fMRI: 13.36 ± 9.50 | 0-63 |
| Morningness-eveningness (MESSi) | Full cohort: 15.96 ± 4.51<br>MSAT: 15.99 ± 4.50<br>task fMRI: 15.21 ± 4.45 | 5-25 |
| Distinctness (MESSi) | Full cohort: 13.55 ± 2.88<br>MSAT: 13.55 ± 2.97<br>task fMRI: 13.40 ± 3.04 | 5-25 |
| Morning affect (MESSi) | Full cohort: 14.82 ± 2.79<br>MSAT: 14.81 ± 2.73<br>task fMRI: 14.45 ± 2.51 | 5-25 |
| Sleep need | Full cohort: 7.85 ± 1.07 <sup>6</sup><br>MSAT: 7.90 ± 1.06 <sup>5</sup><br>task fMRI: 7.85 ± 0.98 <sup>2</sup> | not applicable |
| Sleep quality (PSQI) | Full cohort: 5.73 ± 3.25 <sup>1</sup><br>MSAT: 5.63 ± 3.23 <sup>1</sup><br>task fMRI: 5.92 ± 3.26 <sup>1</sup> | 0-21 |
| Sleep midpoint | Full cohort: 4:13 ± 1:09 <sup>1</sup><br>MSAT: 4:16 ± 1:08 <sup>1</sup><br>task fMRI: 4:03 ± 0:55 <sup>1</sup> | not applicable |
| Sleep debt | Full cohort: 1:06 ± 1:23 <sup>7</sup><br>MSAT: 1:07 ± 1:19 <sup>6</sup><br>task fMRI: 1:06 ± 1:14 <sup>3</sup> | not applicable |

Abbreviations: STAI, State-Trait Anxiety Inventory; BDI, Beck Depression Inventory; MESSi, Morningness-Eveningness-Stability-Scale Improved; PSQI, Pittsburgh Sleep Quality Index.

### 2.2. Moral Sentiment and Action Tendencies task (MSAT)

Participants performed the MSAT task (Zahn et al., 2015; Duan et al., 2023) in the week preceding the fMRI data acquisition. At the beginning of the task, they were asked to provide their name and the name of same-sex best friend that was not their family member or romantic interest. Subsequently, they reported how close they felt with the other person, with the possible scores ranging from 1 (not at all) to 7 (very close). In the current work, we only included data from individuals who reported moderate and strong social bonds (i.e. ≥ 4; n = 140).

During the task participants were faced with hypothetical situations in which either they (self-agency) or their friend (other-agency) were behaving towards one another counter to social and moral values. The paradigm involved 27 identical trials, repeated across the two agency conditions. For each scenario, participants were asked how strongly they would blame themselves and their friend (range: 1-7), what emotion they would most likely feel and what action they would most likely take. The available choices of emotions and actions are presented in **Figure 1**.

**Figure 1.**
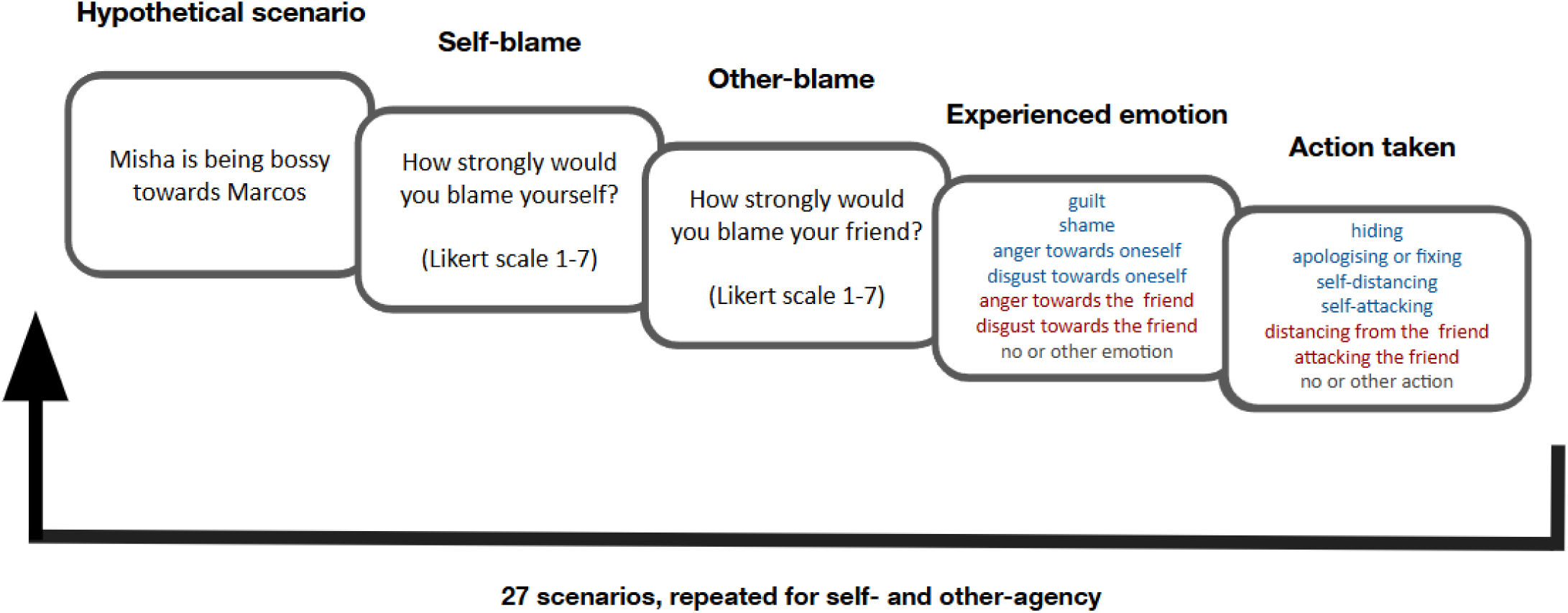
An example trial of the Moral Sentiment and Action Tendencies (MSAT) task. In the paradigm, participants were faced with 27 hypothetical scenarios, repeated across self- and other-agency conditions in which either they or their friend were acting towards the other person counter to social and moral norms. Participants were asked to rate the strength of self- and other-blaming emotions (Likert scale 1-7), as well as the most probably experienced emotion and taken action. Specific self-blaming emotions and related actions are presented in blue, while examples of other-blaming emotions and associated behaviours are shown in red.

### 2.3. Autobiographical guilt recollection fMRI task

#### 2.3.1. Paradigm description

Upon recruitment for the study, participants provided us with 7 instances of autobiographical guilt and neutral memories. For each event, the participants reported the strength of associated guilt feelings and the extent to which they felt they had violated the social norms of behaviour (continuous range: 1-7). For neutral memories, an additional question was asked, probing their emotional valence (continuous range: -3 to 3, i.e. extremely negative to extremely positive). For the subsequent procedures, 5 memories of each type were chosen for every participant, ensuring sufficient guilt ratings for the guilt memories (i.e. ≥4) and low emotional valence for the neutral events (i.e. within the −1 to 1 range).

In the week preceding the fMRI, participants took part in a behavioural experimental session, during which they rated the vividness of emotions associated with each event when recalled outside of the scanner (Likert scale 1-4, i.e. not vividly at all to quite vividly). They also answered questions regarding location, socialness and age of the memories. The provided information was included as a part of the fMRI paradigm, enabling us to confirm the correct retrieval of memories. The described procedures are presented in more detail in the **Supplementary Methods S1.**

The fMRI task was administered using PsychoPy (version 2022.2.5; Peirce et al., 2019) and was presented with MRI-compatible goggles (VisuaStim Digital, Resonance Technology Inc.). The behavioural responses were collected with the right-hand device from the MRI-compatible four-key button box set. Participants were familiarised with the task procedure prior to entering the scanner.

Each trial began with a 10 s presentation of cues pertaining to a specific memory (**Figure 2**). Upon recognising the event, the participants were instructed to ‘relive’ the associated emotions. In the subsequent stage, participants were presented with a question regarding location, socialness or age of the memory (answer limit: 4 s), which was followed by 3 to 5 repetitions of a simple magnitude judgement task (answer limit: 2 s). In the task, being presented with two numbers at the bottom of the screen, the participants had to choose which of them was closer to the target number shown above. The number of arithmetic task repetitions was counterbalanced across the run. Inclusion of the memory-related questions and the magnitude judgement task was meant to shift the attention of scanned individuals from internally- to externally-focused cognition, facilitating easier emotional engagement during subsequent trials. For the same purpose, the guilt and neutral memories were presented in a pseudo-randomised order with not more than two examples of the same condition presented consecutively. Each memory appeared 3 times, every time with a question regarding a different aspect of the event, resulting in 30 trials in total (13 min).

**Figure 2.**
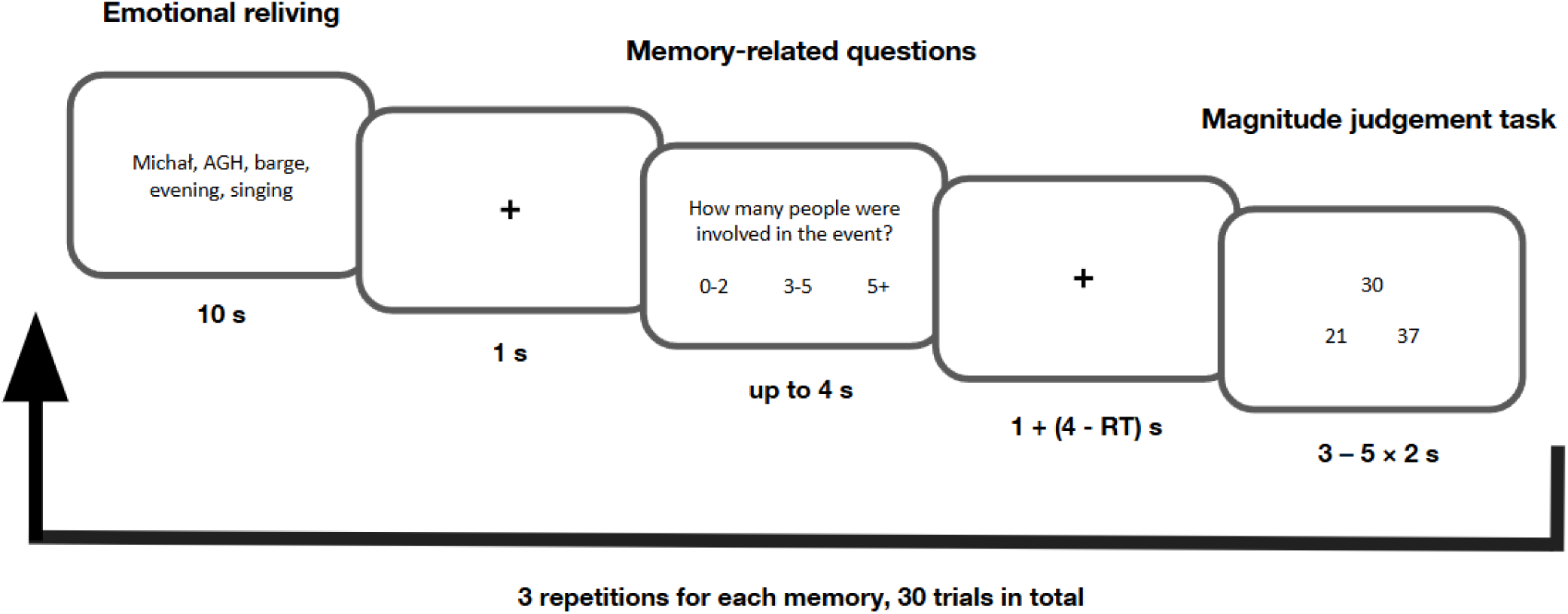
Schematic representation of the autobiographical guilt recollection task. Abbreviations: RT, reaction time.

Following the end of the scanning session, the participants were asked if they had been successful at reliving emotions associated with each memory during all the repetitions, and how vivid the experience had been. The trials for which the participants were unable to perform the task successfully were excluded from the analyses. Last but not least, we additionally probed the approach-avoidance motivation associated with each memory, asking the participants whether they felt like distancing themselves from the involved people, objects and places, or approaching them (continuous range: -3 to 3).

#### 2.3.2. MRI data acquisition

MRI data was acquired with a 3T General Electrics SIGNA Architect scanner. High-resolution structural images were collected with a T1 MP-RAGE sequence. The T2*-weighted fMRI data during autobiographical guilt recollection task was collected using a multiband gradient echo-planar imaging sequence (50 interleaved axial slices; 2.5 mm isotropic voxel; repetition time = 2 s; echo time = 20.4 ms; flip angle = 90°; multiband acceleration factor = 2; left-right phase-encoding direction). In order for signal equilibrium to reach before the beginning of the task, 10 volumes were added at the start of the sequence, resulting in a total fMRI scanning time of 13 min 20 s.

#### 2.3.3. MRI data processing

MRI data was preprocessed with AFNI (Cox, 1996) and FSL (Jenkinson et al., 2012). The anatomical data underwent non-brain tissue removal, warping to the MNI space and segmentation, with the last procedure serving to obtain individual cerebrospinal fluid (CSF) masks. Preprocessing of the functional data began with the removal of the first 5 volumes (non-steady state of the magnetic field), followed by the calculation of motion correction matrix and despiking. Subsequently, the matrices for susceptibility distortion correction and co-registration between anatomical and functional data were obtained. The former procedure was achieved using two separate acquisitions with the opposite phase-encoding directions. All the spatial transformations of the fMRI data were performed in one step, combining the matrices for motion and distortion correction, co-registration and normalisation. The resulting images were smoothed with a 5 mm Gaussian kernel.

The subject-level task activity analysis was performed using AFNI’s *3dREMLfit* program. The nuisance parameters included: motion censoring (separate regressors for each volume with ≥0.5 mm framewise displacement), demeaned original motion parameters and their first-order derivatives (12 parameters in total), the CSF time course (physiological noise) and 6 polynomials to remove the effects of low frequency drifts. The models were additionally run with the pre-whitening option to account for temporal autocorrelations in the data. The brain activity during each part of the trial was modelled separately by convolving box-car responses with the canonical haemodynamic responses. For the emotional reliving epochs, we furthermore included the order of presentation (i.e. first, second or third instance of a particular memory) as a parametric modulator to investigate the habituation effects. The individual brain maps of interest were therefore obtained by contrasting the average activity and habituation effects for emotional reliving of guilt and neutral memories.

In addition to modelling task-evoked activity, we also performed psychophysiological interactions (PPI) analysis using the left (MNI: −50, 4, −10) and right sATL (MNI: 57, −3, −6) as seeds. The regions were defined as 6 mm spheres centered at the peak coordinates from previous self-blame-related neuroimaging studies (Green et al., 2012; Gifuni et al., 2017; Zareba et al., 2024). The physiological regressors were created by extracting seeds’ mean time courses from the residuals of the task activity-related regression, this time run excluding motion censoring to avoid time series discontinuity. Four additional regressors were included, reflecting the average and habituation (parametric modulation) effects during reliving of guilt and neutral memories, compared to the baseline (i.e. physiological effect × psychological effect of interest). The PPI was tested by contrasting the two types of regressors between the emotional and neutral conditions. The regression models additionally included 12 motion parameters, the CSF time course, 6 polynomials and pre-whitening, akin to the analysis probing task-evoked activity.

### 2.4. Statistical analysis

Unless specified otherwise, the described analyses were performed in R (version 4.2.1). The use of permutation-based models was motivated by the non-normal distribution of the data.

#### 2.4.1. Sleep and mental health psychometric measures

Permutation-based linear models (10000 iterations) testing whether sleep and mental health measures were associated with chronotype, distinctness and their interaction were run using the *lmperm* function from the *permuco* library (Frossard and Renaud, 2021). Age and sex were controlled as covariates. False discovery rate (FDR < 0.05) was used to correct for multiple comparisons, controlling for the number of tested metrics.

#### 2.4.2. MSAT

The strength of self- and other-blaming emotions was averaged within self- and other-agency conditions. Similarly, we calculated per condition the proportion of trials in which participants chose particular emotions and actions. Such data served as the dependent variables for permutation-based repeated measures analyses of covariance (ANCOVAs) run using the *aovperm* function from the *permuco* library (Frossard and Renaud, 2021) with 10000 permutations. The models tested the main effects and interactions between chronotype, distinctness and agency (within-subject condition), controlling for age and sex. FDR was applied separately for each metric category, i.e. strength of self- and other-blaming emotions, experienced emotions and reported actions.

#### 2.4.3. Autobiographical guilt recollection task neuroimaging data

The autobiographical guilt recollection task was performed by 80 individuals. To avoid a confounding influence of partial sleep restriction, we only included participants whose neuroimaging sessions began at least 1 hour after their habitual wake up time, excluding 5 individuals. To study the habituation effects, we furthermore included only those individuals who reported successful reliving of guilt and neutral memories for at least 3 out of 5 trials across each repetition, excluding 2 participants. The final sample therefore consisted of 73 individuals.

The neuroimaging data was analysed in AFNI and R. Separate models were run for the average and habituation effects for task-evoked activity and task-dependent sATL functional connectivity. TOD when individuals performed the task was derived from the raw neuroimaging files (range: 8:31 – 21:45). The models investigated the main effects and interactions of individual chronotype, distinctness and TOD, controlling for age and sex.

The whole-brain calculations were run with AFNI’s 3dMVM program (Chen et al., 2014). The results were controlled for multiple comparisons at the cluster-level using family-wise error rate correction (FWE < 0.05) following voxel-level thresholding at p < 0.001. Additionally, to study the sATL - sgACC circuitry in greater detail, we deployed a region-of-interest (ROI)-based approach using R’s *lm* function. The left and right sATL ROIs were delineated as described previously (section 2.3.3). The bilateral sgACC region was defined as a 6 mm sphere centered at the following MNI coordinates (X, Y, Z): 0, 15, -5 (Moll et al., 2006; Zareba et al., 2024). The measures of interest included each region’s mean brain activity, and the PPI of the left and right sATL with sgACC. Multiple comparison correction was achieved using FDR.

#### 2.4.4. Autobiographical guilt recollection task behavioural data

Mean values were calculated for the following metrics collected for both guilt and neutral memories: strength of guilt feelings, perceived social code violation, vividness of memory recall both outside and inside the scanner, approach–avoidance motivation, salience (defined as the absolute value of the approach–avoidance dimension), and the accuracy and reaction times (RTs) for the memory-related questions. We also calculated the total number of memories of each type that participants were able to successfully relive in the scanner. For neutral memories, an additional metric was their mean emotional valence. Furthermore, individual differences in the approach-avoidance motivation and salience between guilt and neutral memories were computed. For the magnitude judgment task, accuracy and mean RTs were obtained. Due to the use of incorrect buttons, one and three participants were excluded from the analysis of RTs and accuracy for, respectively, the memory- and arithmetics-related parts of the task.

The statistical analysis was performed using *permuco*’s *lmperm* function (10000 permutations; Frossard and Renaud, 2021). For the measures collected before the neuroimaging session, the models tested whether they were associated with the main effects and interaction of chronotype and distinctness, controlling for the effects of age and sex. In the case of data obtained during the fMRI session, the respective models additionally included the main effect of TOD, together with its two- and three-way interactions with chronotype and distinctness. FDR was applied separately for the data pertaining to the guilt and neutral memories, the differences between the two, and the magnitude judgement task, controlling for the number of models in which a particular term appeared.

## 3. Results

### 3.1. Associations of chronotype and distinctness with sleep and mental health measures

Higher eveningness was associated with later sleep midpoint (T = 4.91, p_FDR_ < 0.001), higher sleep debt (T = 2.54; p_FDR_ = 0.025), decreased morning affect (T = -4.59, p_FDR_ < 0.001) and elevated trait-anxiety (T = 2.18, p_FDR_ = 0.049). Nominally significant moderation effects of distinctness were observed, where higher distinctness strengthened the links of eveningness with trait-anxiety (T = 2.14; p_uncorr._ = 0.035; p_FDR_ = 0.147) and sleep debt (T = 2.02; p_uncorr._ = 0.042; p_FDR_ = 0.147; **Supplementary Figure S1**). On the nominal level, higher eveningness (T = 2.02; p_uncorr._ = 0.045; p_FDR_ = 0.063) and distinctness (T = 2.22; p_uncorr._ = 0.028; p_FDR_ = 0.193) were also independently related to higher reported sleep need. The full results, including the non-significant associations with subjective sleep quality and depressive symptomatology, are described in **Supplementary Table S1**.

### 3.2. Associations of chronotype and distinctness with self- and other-blaming emotions and action tendencies

The analysis of the strength of self- and other-blaming emotions in the MSAT task revealed significant interactions between chronotype and agency. Higher eveningness was related to less pronounced self-blaming in the self-agency condition and increased self-blaming in the other-agency trials (F = 5.25; p_FDR_ = 0.039; **Figure 3A**). The opposite association was seen for the other-blaming emotions: they were increased in the self-agency trials and decreased in the other-agency condition (F = 4.28; p_FDR_ = 0.039; **Figure 3B**). With regards to action tendencies, later chronotype was related to reduced self-distancing (F = 7.91; p_FDR_ = 0.041, **Figure 3C**). Additionally, a nominally significant interaction of chronotype and agency was found, driven predominantly by decreased distancing from others in the other-agency condition in evening types (F = 5.34; p_uncorr._ = 0.023; p_FDR_ = 0.162; **Figure 3D**). The nominally significant associations of distinctness with action tendencies (p_uncorr._ < 0.05) are shown in **Supplementary Figure S2**. No links with the frequency of specific self- and other-blaming emotions were found for either chronotype or distinctness. Full results are presented in **Supplementary Tables S2-3.**

**Figure 3.**
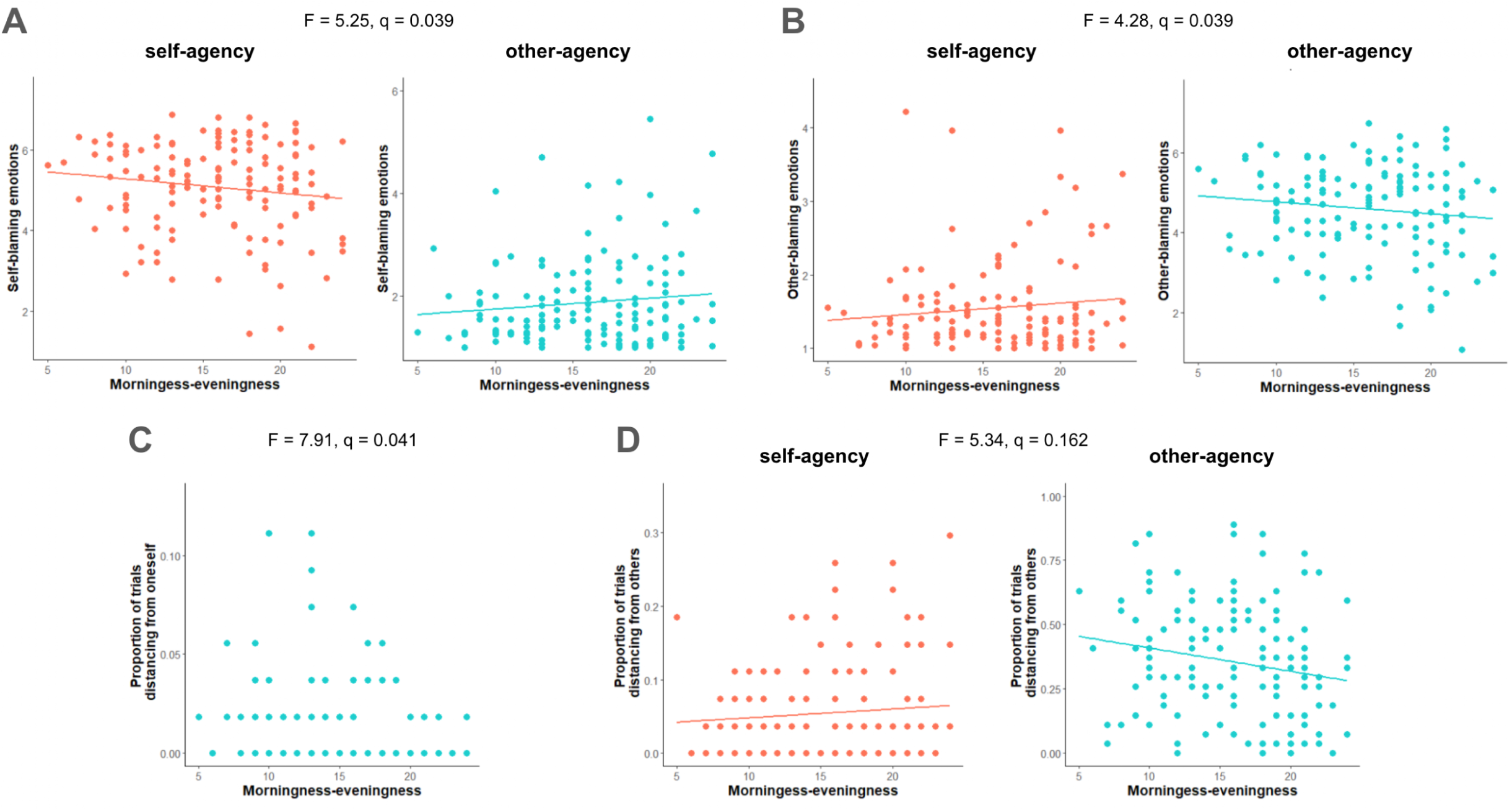
Associations of morningness-eveningness with self-blaming emotions and action tendencies from the Moral Sentiment and Action Tendencies Task (MSAT). Greater eveningness was associated with (A) weaker self-blaming and (B) stronger other-blaming emotions in the self-agency condition. An opposite pattern was observed in the other-agency trials (A-B). Later chronotype was furthermore associated with distancing behaviours: (C) reduced self-distancing, regardless of the condition, together with (D) increased and decreased distancing from others in the self- and other-agency trials, respectively (trend-level). Abbreviations: q, false discovery rate-corrected p value.

### 3.3. Associations of diurnal rhythmicity with neural correlates of reliving autobiographical guilt memories

The whole brain models testing the average task effects revealed only a single significant result, i.e. an interaction of distinctness and TOD for the task-related connectivity between the right sATL and the left hippocampus (**Table 2**). In individuals with higher subjective diurnal rhythm amplitude, lower coupling between the two regions was observed at later TOD (**Figure 4**). The opposite association was present in participants with lower distinctness.

**Table 2.** Summary of the brain activity and connectivity patterns associated with diurnal rhythmicity. For self-blame-dependent functional connectivity results, the first area represents the seed region. The multiplication sign represents interaction effects.

| Effect type | Neural measure | Brain region | MNI | Voxels | Statistics |
| --- | --- | --- | --- | --- | --- |
| ***Morningness-eveningness*** |  |  |  |  |  |
| Average | Task activity | B subgenual anterior cingulate cortex | 0, 15, -5 | 61 | T = -2.79<sup>R</sup> |
| Average | Task activity | R superior anterior temporal lobe | 57, -3, -6 | 55 | T = -2.56<sup>R</sup> |
| ***Distinctness*** |  |  |  |  |  |
| Habituation | Task-dependent connectivity | R superior anterior temporal lobe<br>L cerebellar lobule III<br>L parabrachial nuclei<sup>1</sup> | 57, -3, -6<br>-14, -38, -28 | 55<br>22 | F = 30.23<sup>W</sup> |
| ***Time-of-day*** |  |  |  |  |  |
| Habituation | Task activity | B precuneus | -2, -60, 32 | 44 | F = 20.44<sup>W</sup> |
| ***Distinctness × time-of-day*** |  |  |  |  |  |
| Average | Task-dependent connectivity | R superior anterior temporal lobe<br>L hippocampus | 57, -3, -6<br>-24, -18, -12 | 55<br>41 | F = 27.13<sup>W</sup> |
| ***Morningness-eveningness***<br>***× distinctness × time-of-day*** |  |  |  |  |  |
| Average | Task-dependent connectivity | R superior anterior temporal lobe<br>B subgenual anterior cingulate cortex | 57, -3, -6<br>0, 15, -5 | 55<br>61 | T = 2.72<sup>R</sup> |

Abbreviations: L, left; R, right; B, bilateral; <sup>R</sup>, region-of-interest analysis, $p_{FDR} < 0.05$ ; <sup>W</sup>, whole-brain analysis, cluster-level $p_{FWE} < 0.05$ .
<sup>1</sup> The location was confirmed using a high resolution human brainstem atlas (Bianciardi et al., 2016; see **Supplementary Figure S3**).

**Figure 4.**
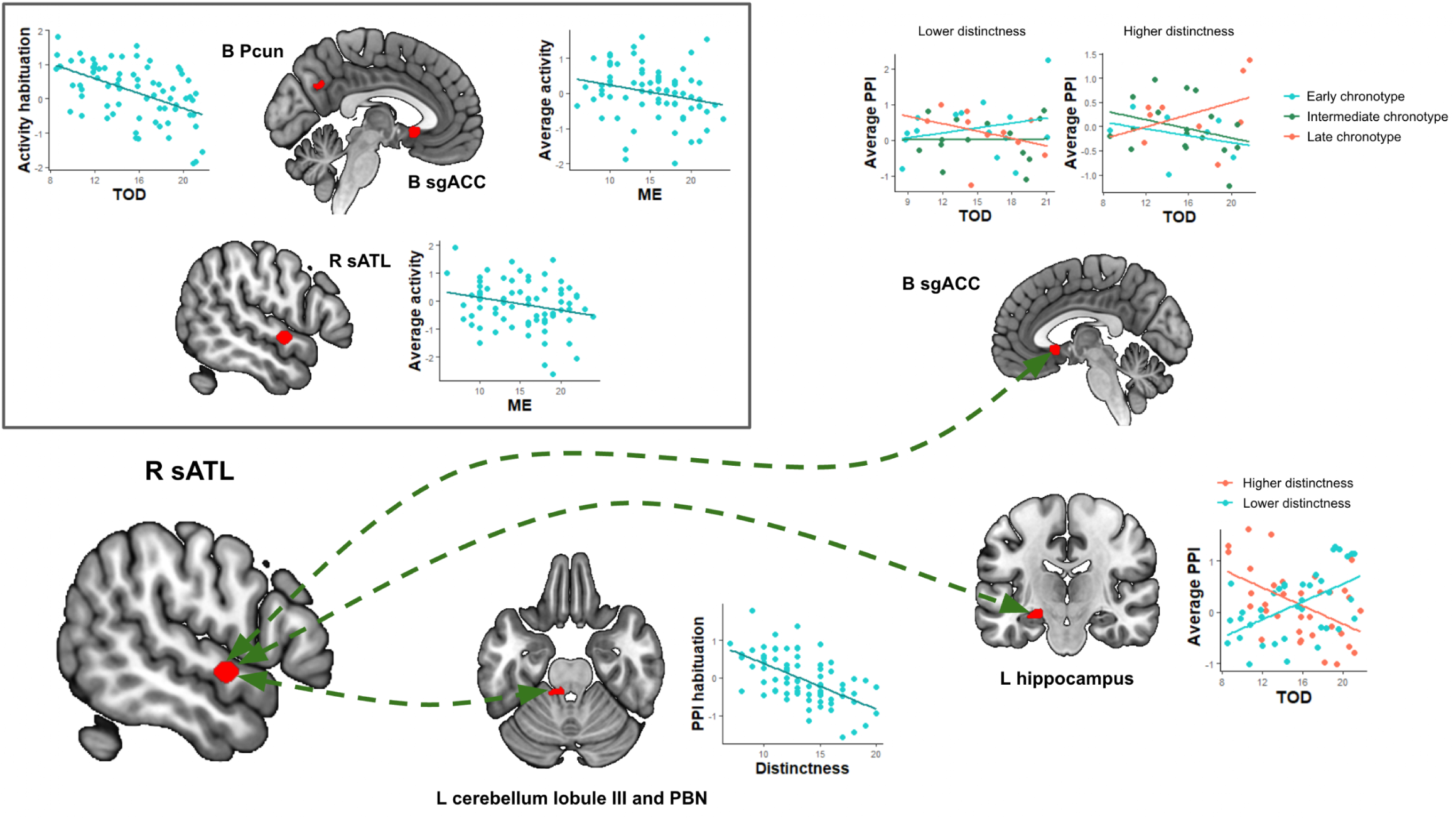
Summary of the neuroimaging results. The findings associated with differences in brain activity are shown in the box in the top left corner. The results pertaining to the task-related functional connectivity are presented using green lines connecting the seed region (i.e. the right superior anterior temporal lobe; R sATL) with the brain areas where significant effects of diurnal measures were found. For the visualisation of the interaction effects, the cohort was stratified into multiple distinctness (lower and higher) and chronotype groups (early, intermediate and late). The details are presented in **Supplementary Methods S2**. Abbreviations: TOD, time-of-day; ME, morningness-eveningness; PPI, psychophysiological interaction; B, bilateral; L, left; R, right; Pcun, precuneus; sgACC, subgenual anterior cingulate cortex; sATL, superior anterior temporal lobe; PBN, parabrachial nuclei.

Meanwhile, the ROI analysis focused specifically on the sATL-sgACC circuitry revealed that eveningness was negatively related to self-blaming-evoked activity in the bilateral sgACC (T = -2.79, β = -0.061, p_FDR_ = 0.019) and the right sATL (T = -2.56, β = -0.059, p_FDR_ = 0.019). Additionally, a three-way interaction between chronotype, distinctness and TOD was observed for the task connectivity between the right sATL and bilateral sgACC (T = 2.72, β = 0.004, p_FDR_ = 0.017). For participants with higher distinctness, stronger coupling between the regions was observed at individually preferred TOD, i.e. morning for morning types and evening for evening types (**Figure 4**). The effects appeared reversed in participants with lower subjective diurnal amplitude.

With respect to the habituation analyses, the whole-brain models revealed stronger decreases in the bilateral precuneus activity across the guilt memory repetitions at later TOD (**Table 2**). Stronger habituation was also observed for the coupling of the right sATL with the left cerebellar lobule III and parabrachial nuclei in individuals with greater distinctness. The ROI analyses yielded no significant habituation-related findings within the sATL-sgACC circuitry.

The significant neuroimaging findings are summarised in **Table 2** and **Figure 4**. The full results of the ROI analyses can be found in **Supplementary Tables S4-5.**

### 3.4. Associations of diurnal rhythmicity with behavioural measures of the autobiographical guilt recollection task

A single association significant at the FDR-corrected level was found, i.e. greater difference in the salience between the guilt and neutral memories for participants who performed the task at later TOD (T = 2.35, p_FDR_ = 0.049; **Figure 5A**). On the nominal level, a three-way interaction between chronotype, distinctness and TOD was observed (T = 2.00, p_uncorr._ = 0.049, p_FDR_ = 0.293; **Figure 5B**), where individuals with higher distinctness displayed increased avoidance towards their guilt memories at non-preferred TOD. The effects appeared reversed and diminished in participants with lower subjective diurnal amplitude.

**Figure 5.**
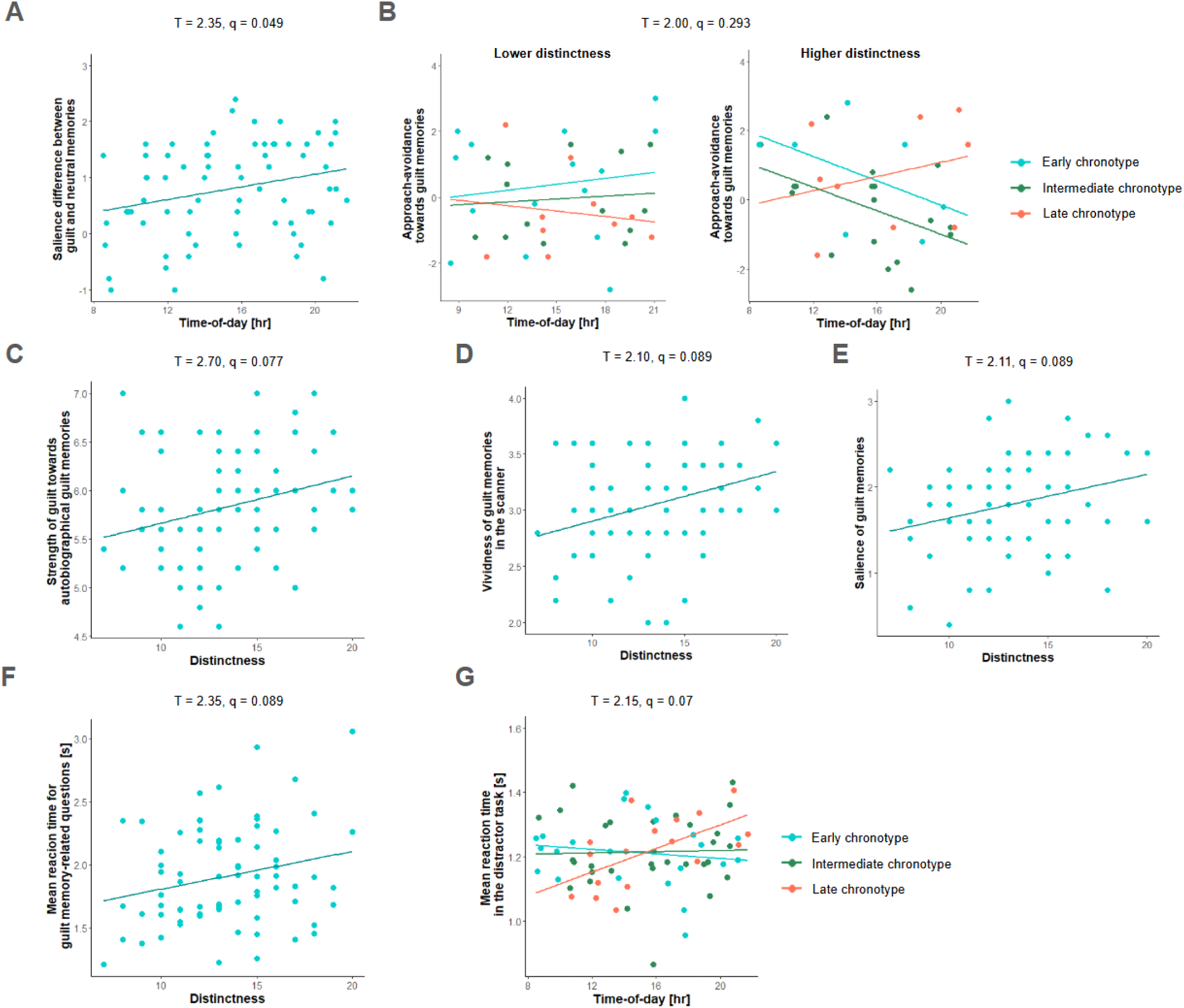
Significant (A) and trend-level (B-G) associations between diurnal measures or their interactions and behavioural data derived from the autobiographical guilt recollection task. For the visualisation of the interaction effects, the cohort was stratified into multiple distinctness (lower and higher) and chronotype groups (early, intermediate and late). The details are presented in **Supplementary Methods S2**. Abbreviations: q, false discovery rate-corrected p value.

Additionally, we observed trend-level links between distinctness and the following characteristics of the guilt memories (**Figure 5C-F)**: strength of guilt feelings (T = 2.70, p_uncorr._ = 0.009, p_FDR_ = 0.077), vividness of the in-scanner recall (T = 2.10, p_uncorr._ = 0.039, p_FDR_ = 0.089), salience (T = 2.11, p_uncorr._ = 0.039, p_FDR_ = 0.089) and mean RT for the memory content-related questions (T = 2.35, p_uncorr._ = 0.022, p_FDR_ = 0.089). Last but not least, an interaction between TOD and chronotype was observed for the mean RT in the distractor task, driven by higher values in eveningness-oriented individuals at later TOD (T = 2.15, p_uncorr._ = 0.035, p_FDR_ = 0.07; **Figure 5G**). The full results are shown in **Supplementary Table S6.**

## 4. Discussion

The current study confirms the previously reported associations of eveningness with adverse affective (i.e. trait-anxiety) and sleep health (i.e. sleep debt) outcomes, hinting that these effects may be more pronounced in individuals with higher distinctness (i.e., trend-level). Simultaneously, in the MSAT task, late chronotypes demonstrated diminished self-distancing and contextually-maladaptive blaming, i.e. decreased self-blaming and increased other-blaming emotions in the self-agency condition. This pattern was reversed for the other-agency trials. Last but not least, the neuroimaging task revealed that diurnal rhythmicity modulated neural and behavioural characteristics of reliving autobiographical guilt memories, providing plausible correlates of adverse emotional health outcomes in late chronotypes. Importantly, while some of the findings appear to have trait-like characteristics (i.e. main effects of chronotype and distinctness), some others may reflect state-like processes changing with TOD, underlining complex associations between diurnal rhythmicity and affective functioning.

One of the novel aspects of our work considered how interactions between chronotype and distinctness affected sleep and mental health measures. We observed no modulatory effects of distinctness on core chronobiological features of chronotype, i.e. sleep midpoint and morning affect. On the trend-level, however, distinctness strengthened the associations of eveningness with adverse health outcomes, i.e. trait-anxiety and sleep debt. Our results suggest that the sleep debt-related findings might be partially related to increased sleep need of late chronotype and high distinctness individuals, raising a possibility that both dimensions of diurnal rhythms could be associated with differences in homeostatic sleep pressure regulation (Mongrain and Dumont, 2007). An important caveat of this cross-sectional work is, however, that we cannot disentagle the bidirectional associations between sleep and anxiety (Ariño-Braña et al., 2025). Larger prospective studies, encompassing broader age ranges, are therefore needed to complement these preliminary findings.

One pathway through which increased anxiety may manifest in late chronotypes leads through self-blaming emotions and associated behaviours. Indeed, evening-types exhibited reduced self-distancing, consistent with our previous finding of diminished self-distancing in anxious individuals experiencing shame or self-anger (Zareba et al., 2026). This propensity to focus on self-centered thoughts in social scenarios might lead to ruminations, a characteristic of affective disorders that has been previously reported in late chronotypes (Antypa et al., 2017). The finding on the coping strategies was paralleled by apparent misattribution of blame, i.e. increased blaming of the action recipients rather than agents. On one hand, stronger self-blaming in the other-agency condition could be potentially explained by increased sense of responsibility for actions of others in anxious individuals (Apetroaia et al., 2015). Simultaneously, when experiencing negative emotions, individuals with emotion regulation difficulties are more likely to ascribe to others the responsibility for their own actions (Kaufmann et al., 2022). The presented line of interpretation remains, nonetheless, speculative and specific cognitive mechanisms behind the reported results should be further investigated.

Complementing the behavioural findings, neuroimaging data showed that late chronotypes had decreased activity in key self-blame-related regions, i.e. bilateral sgACC and right sATL. The sgACC has been associated with perceived group belongingness (Rüsch et al., 2014) and prediction errors during social feedback affecting self-esteem (Will et al., 2017). Its decreased activity during guilt reliving may therefore reflect greater feelings of social exclusion following transgressions. Consistently, loneliness has been shown to mediate the relationship between eveningness and social anxiety symptoms (Zhu et al., 2024). The role of sATL in social semantic processing may likewise contribute to negative cognitive biases observed in late chronotypes (Thye et al., 2024). For instance, akin to affective disorders patients, they show enhanced recognition of negative facial expressions (Horne et al., 2016). Decreased sATL activity in related paradigms has previously been associated with negative affectivity (Kret et al., 2011), suggesting that cognitive biases might be related to more efficient access to these specific emotional semantic representations. This aligns with semantic models suggesting that retrieval efficiency increases when a concept is accessed more frequently as it becomes more established within the semantic network, thereby lowering metabolic demands in the ATL (Lambon Ralph et al. 2017; Rogers et al. 2004). Distinctness, in turn, was associated with stronger habituation of right sATL connectivity with cerebellar lobule III and parabrachial nuclei, possibly reflecting proprioceptive processing following guilt-induced autonomic activity (Schmahmann, 2018; Palmiter, 2018; Golde et al., 2023; Zareba et al., 2026). Together with previous findings, this suggests that distinctness may primarily modulate subcortical and lower-order cortical regions (Zareba et al., 2023; Scislewska et al., 2025).

The neural patterns described above were independent of TOD and may therefore represent trait-like effects. Importantly, our findings suggest that affective brain processing reflects an interplay between such stable characteristics and state-like processes varying with TOD. For example, stronger evening habituation of precuneus activity may indicate facilitated guilt memory retrieval across epochs (Flanagin et al., 2023), potentially reflecting diurnal differences in subjective memory characteristics. Nonetheless, the TOD-related difference in salience between guilt and neutral memories was correlated with the precuneus habituation effect only at the trend-level (r = -0.21; p = 0.071), indicating that other factors are at play. State-like effects were also observed for the right sATL connectivity: two-way interactions of TOD with distinctness and three-way interactions with distinctness and chronotype emerged in the left hippocampus and bilateral sgACC, respectively. Hippocampus contributes to processing of temporal aspects and detailed scenic reconstruction of episodic memories (Flanagin et al., 2023). As such, our work suggests that the interactions between episodic and semantic memory, including processes like generalisation, might depend on individual diurnal rhythmicity (Rolls et al., 2025). Similarly, diurnal modulation of the sATL - sgACC coupling, a putative biomarker of integrating social conceptual information with self-worth and affiliative processing (Zahn, 2025), raises important implications for its behavioural relevance. As increased self-blame-dependent connectivity between these regions is observed across clinically depressed and subclinically anxious populations (Lythe et al., 2015; Zareba et al., 2026), individuals may be more susceptible to the negative effects of self-blame on self-esteem at TODs when coupling is the strongest. If confirmed, such findings may open avenues for the development of psychological treatments tailored to individual diurnal rhythmicity.

### Limitations

An important caveat of the state-like neuroimaging results is, however, the cross-sectional character of the study. Prior to translational applications, the presented findings should be replicated using within-subject designs. Furthermore, individual diurnal rhythmicity was tested with self-reports. Although the morningness-eveningness scale has been validated against objective circadian measures, such as dim-light melatonin onset (Kantermann et al., 2015), relatively little is known about the molecular correlates of the distinctness dimension. Future studies are encouraged to fill this gap. We additionally note the use of clock time and not individual circadian time (i.e. hours after wake), and the fact that the current work was conducted in a naturalistic cohort performing their daily routines. This prevented clear delineation between circadian and homeostatic sleep factors (Cajochen and Schmidt, 2025) but, nonetheless, increased the ecological validity of the results.

## 5. Conclusions

In summary, the presented findings extend the links between late chronotype and emotional dysregulation by demonstrating maladaptive self-blaming behaviours and altered activity in the self-blaming circuitry in that group. Additionally, they suggest that affective processing reflects an interplay between stable trait-like characteristics and dynamic state-like processes changing across the day, contributing to fine-grained understanding of how emotional cognition is embedded in the brain and encouraging development of chronobiology-informed treatments.

## Data availability

The guilt recollection task is available on the following GitHub repository: https://github.com/mrzareba/10.1016-j.pnpbp.2026.111679/. Maps of mean tSNR and average task-evoked activity for the guilt recollection task in a larger cohort (n = 80) are available as a part of the following NeuroVault repository: https://neurovault.org/collections/21877/. Individual statistical maps generated for the current study may be shared upon request.

## CRediT author statement

MRZ: Conceptualization, Methodology, Investigation, Formal analysis, Writing - Original Draft, Visualisation. IGG: Resources, Investigation, Writing - Review & Editing. MIM: Resources, Investigation, Writing - Review & Editing. MV: Supervision, Conceptualisation, Methodology, Investigation, Writing - Review & Editing.

## Ethical statement

All the procedures followed were in accordance with the ethical standards of the responsible committee on human experimentation (institutional and national) and with the Declaration of Helsinki (1975), and the applicable revisions at the time that this research was underway. Informed consent to be included in the study was obtained from all the participants. The study protocol was approved by the Universitat Jaume I Ethics Committee (CEISH/07/2022).

