## Supplementary Material for "Diurnal rhythmicity modulates self-blaming behaviours and neural processing of autobiographical guilt memories in the trait-like and state-like manner"

### Supplementary Methods S1

Upon recruitment for the study, participants provided us with 7 instances of autobiographical guilt and neutral memories. Ensuring participants' comfort, for each memory, they provided 5 cues, making the exact content of personal events unknown to the experimenters. To make the cues comparable within and across the individuals, we required at least one cue pertaining to the involved people, an associated abstract or concrete object and the location of the event. In some cases the exact wording of the cues was slightly altered by the experimenters for brevity, and so the participants were later asked to confirm their ability to correctly identify the memories.

At this initial stage, the participants additionally provided us with information on how guilty they felt about the reported events, together with the extent to which they felt they had violated the social norms of behaviour (continuous range: 1-7). In the case of neutral events, we further asked participants about their emotional valence (continuous range: -3 to 3, i.e. extremely negative to extremely positive). Five memories of each type were chosen for every participant. The subsequent parts of the experiment were performed only by individuals reporting guilt ratings ≥4 for the guilt memories, and emotional valence ratings within the −1 to 1 range for the neutral memories.

In the week preceding the fMRI, participants took part in a behavioural experimental session, during which they rated how vividly they could recall outside the scanner the emotions associated with each event (Likert scale 1-4, i.e. not vividly at all to quite vividly). For each memory, they were additionally asked to respond to three more questions with prespecified answers concerning the type of location, number of people involved and the time that had passed since each event took place. The provided information was included as a part of the paradigm performed inside the scanner, enabling us to confirm the correct retrieval of memories.

### Supplementary Methods S2

For the visualisation of the interaction effects in **Figures 4** and **5**, the cohort was stratified into multiple distinctness (lower and higher) and chronotype groups (early, intermediate, late). The lower distinctness group encompasses individuals with distinctness values ranging from 7 to 13 (maximum value: 25), and the higher distinctness group includes participants with scores from 14 to 20. Consequently, the early chronotype group consists of individuals scoring 6 to 12 on the morningness-eveningness scale (maximum value: 25), the intermediate chronotype group includes participants with scores from 13 to 18, and the late chronotype group encompasses subjects with the values between 19 and 24.

**Supplementary Table S1.** The main and interaction effects of morningness-eveningness and distinctness on sleep and mental health measures. The results surviving the multiple comparison correction ($p_{FDR} < 0.05$) are shown in blue and bold, while nominally significant findings ($p_{uncorr.} < 0.05$) are additionally presented in italics.

| Variable | Morningness-eveningness | | Distinctness | | Morningness-eveningness × Distinctness | |
|---|---|---|---|---|---|---|
| | **T-stat** | **$p_{uncorr.}$ ($p_{FDR}$)** | **T-stat** | **$p_{uncorr.}$ ($p_{FDR}$)** | **T-stat** | **$p_{uncorr.}$ ($p_{FDR}$)** |
| Trait-anxiety | **2.18** | **0.028 (0.049)** | 0.75 | 0.455 (0.648) | ***2.14*** | ***0.035 (0.147)*** |
| Depressive symptoms | 1.71 | 0.086 (0.100) | 0.24 | 0.810 (0.870) | 1.07 | 0.289 (0.405) |
| Morning affect | **-4.59** | **<0.001 (<0.001)** | 1.62 | 0.108 (0.262) | -1.50 | 0.137 (0.320) |
| Sleep need | ***2.02*** | ***0.045 (0.063)*** | ***2.22*** | ***0.028 (0.193)*** | 1.13 | 0.263 (0.405) |
| Sleep quality | 1.18 | 0.241 (0.241) | 0.74 | 0.463 (0.648) | 0.51 | 0.612 (0.714) |
| Sleep debt | **2.64** | **0.011 (0.025)** | 1.61 | 0.112 (0.262) | ***2.02*** | ***0.042 (0.147)*** |
| Sleep midpoint | **4.91** | **0.001 (<0.001)** | -0.17 | 0.870 (0.870) | 0.27 | 0.783 (0.783) |

Abbreviations: FDR, false discovery rate.

**Supplementary Table 2.** The effects of morningness-eveningness, distinctness and their interactions with each other and agency on the self- and other-blaming emotions in the Moral Sentiment and Action Tendencies (MSAT) task. The results surviving the multiple comparison correction ($p_{FDR} < 0.05$) are shown in blue and bold.

| Variable | Morningness-eveningness | | Distinctness | | Morningness-eveningness × distinctness | | Morningness-eveningness × agency | | Distinctness × agency | | Morningness-eveningness × distinctness × agency | |
|---|---|---|---|---|---|---|---|---|---|---|---|---|
| | F-stat | $p_{uncorr}$ ($p_{FDR}$) | F-stat | $p_{uncorr}$ ($p_{FDR}$) | F-stat | $p_{uncorr}$ ($p_{FDR}$) | F-stat | $p_{uncorr}$ ($p_{FDR}$) | F-stat | $p_{uncorr}$ ($p_{FDR}$) | F-stat | $p_{uncorr}$ ($p_{FDR}$) |
| **Self-blaming emotions strength** | 0.34 | 0.56 (0.56) | 1.94 | 0.164 (0.328) | 0.94 | 0.338 (0.596) | **5.25** | **0.021 (0.039)** | 0.14 | 0.705 (0.705) | 0.65 | 0.414 (0.668) |
| **Other-blaming emotions strength** | 0.74 | 0.393 (0.56) | 0.90 | 0.352 (0.352) | 0.27 | 0.596 (0.596) | **4.28** | **0.039 (0.039)** | 0.60 | 0.439 (0.705) | 0.19 | 0.668 (0.668) |
| **Shame** | 0.20 | 0.662 (0.927) | 1.34 | 0.243 (0.789) | 0.01 | 0.990 (0.990) | 0.62 | 0.433 (0.690) | 0.04 | 0.835 (0.835) | 2.56 | 0.115 (0.632) |
| **Guilt** | 2.00 | 0.165 (0.816) | 0.02 | 0.899 (0.991) | 0.50 | 0.476 (0.703) | 2.48 | 0.121 (0.467) | 0.26 | 0.607 (0.835) | 0.40 | 0.537 (0.632) |
| **Anger towards oneself** | 0.03 | 0.867 (0.967) | 0.92 | 0.338 (0.789) | 1.04 | 0.310 (0.703) | 0.43 | 0.510 (0.690) | 2.60 | 0.109 (0.536) | 0.82 | 0.365 (0.632) |
| **Contempt towards oneself** | 0.53 | 0.466 (0.816) | 0.09 | 0.768 (0.991) | 0.01 | 0.918 (0.990) | 0.03 | 0.853 (0.853) | 0.09 | 0.755 (0.835) | 0.02 | 0.906 (0.906) |
| **Anger towards the friend** | 0.76 | 0.389 (0.816) | 1.30 | 0.252 (0.789) | 1.35 | 0.247 (0.703) | 0.29 | 0.591 (0.690) | 2.09 | 0.153 (0.536) | 1.66 | 0.196 (0.632) |
| **Contempt towards the friend** | 0.00 | 0.967 (0.967) | 0.02 | 0.887 (0.991) | 0.45 | 0.502 (0.703) | 1.63 | 0.210 (0.490) | 0.14 | 0.717 (0.835) | 0.47 | 0.497 (0.632) |
| **No or other emotion** | 0.57 | 0.450 (0.816) | 0.00 | 0.991 (0.991) | 1.81 | 0.179 (0.703) | 2.30 | 0.133 (0.466) | 0.07 | 0.798 (0.835) | 0.37 | 0.542 (0.632) |

Abbreviations: FDR, false discovery rate.

**Supplementary Table 3.** The effects of morningness-eveningness, distinctness and their interactions with each other and agency on the action tendencies in the Moral Sentiment and Action Tendencies (MSAT) task. The results surviving the multiple comparison correction ($p_{FDR} < 0.05$) are shown in blue and bold, while nominally significant findings ($p_{uncorr.} < 0.05$) are additionally presented in italics.

| **Variable** | **Morningness-eveningness** | | **Distinctness** | | **Morningness-eveningness × distinctness** | | **Morningness-eveningness × agency** | | **Distinctness × agency** | | **Morningness-eveningness × distinctness × agency** | |
|---|---|---|---|---|---|---|---|---|---|---|---|---|
| | **F-stat** | **$p_{uncorr}$ ($p_{FDR}$)** | **F-stat** | **$p_{uncorr}$ ($p_{FDR}$)** | **F-stat** | **$p_{uncorr}$ ($p_{FDR}$)** | **F-stat** | **$p_{uncorr}$ ($p_{FDR}$)** | **F-stat** | **$p_{uncorr}$ ($p_{FDR}$)** | **F-stat** | **$p_{uncorr}$ ($p_{FDR}$)** |
| **Distancing from oneself** | **7.91** | **0.006 (0.041)** | 0.41 | 0.531 (0.925) | 0.99 | 0.318 (0.959) | 2.22 | 0.133 (0.331) | 0.34 | 0.567 (0.992) | 0.26 | 0.609 (0.836) |
| **Attacking oneself** | 0.00 | 0.996 (0.996) | 2.05 | 0.158 (0.716) | 0.48 | 0.463 (0.959) | 0.32 | 0.582 (0.679) | 0.03 | 0.856 (0.999) | 1.35 | 0.231 (0.609) |
| **Hiding** | 0.30 | 0.587 (0.685) | 0.07 | 0.796 (0.925) | 0.01 | 0.970 (0.970) | 0.09 | 0.762 (0.762) | 0.08 | 0.769 (0.999) | 1.23 | 0.261 (0.609) |
| **Apologising or fixing** | 1.15 | 0.293 (0.685) | 1.56 | 0.217 (0.716) | 0.03 | 0.872 (0.970) | 0.31 | 0.576 (0.679) | ***6.37*** | ***0.014 (0.095)*** | 0.04 | 0.836 (0.836) |
| **Attacking the friend** | 0.61 | 0.435 (0.685) | 0.17 | 0.678 (0.925) | 0.04 | 0.837 (0.970) | 0.87 | 0.360 (0.630) | 1.03 | 0.314 (0.733) | 0.08 | 0.774 (0.836) |
| **Distancing from the friend** | 0.61 | 0.422 (0.685) | 1.03 | 0.307 (0.716) | 1.34 | 0.249 (0.959) | ***5.34*** | ***0.023 (0.162)*** | 0.00 | 0.999 (0.999) | 0.30 | 0.580 (0.836) |
| **No or other action** | 0.30 | 0.583 (0.685) | 0.00 | 0.925 (0.925) | 0.34 | 0.548 (0.959) | 2.18 | 0.142 (0.331) | ***4.60*** | ***0.036 (0.126)*** | 2.22 | 0.132 (0.609) |

Abbreviations: FDR, false discovery rate.

**Supplementary Table 4.** The main and interactions effects of morningness-eveningness, distinctness and time-of-day on average task-evoked activity and connectivity. The results surviving the multiple comparison correction ($p_{FDR} < 0.05$) are shown in blue and bold, while nominally significant findings ($p_{uncorr.} < 0.05$) are additionally presented in italics.

| Neural measure | Brain region | T-stat | β (SE) | $p_{uncorr.}$ ($p_{FDR}$) |
|---|---|---|---|---|
| ***Morningness-eveningness*** | | | | |
| Task activity | L superior anterior temporal lobe | -1.14 | -0.025 (0.022) | 0.259 (0.259) |
| **Task activity** | **R superior anterior temporal lobe** | **-2.56** | **-0.059 (0.023)** | **0.013 (0.019)** |
| **Task activity** | **B subgenual anterior cingulate cortex** | **-2.79** | **-0.061 (0.022)** | **0.007 (0.019)** |
| Task-dependent connectivity | L superior anterior temporal lobe<br>B subgenual anterior cingulate cortex | 1.09 | 0.020 (0.019) | 0.282 (0.564) |
| Task-dependent connectivity | R superior anterior temporal lobe<br>B subgenual anterior cingulate cortex | 0.03 | 0.001 (0.019) | 0.978 (0.978) |
| ***Distinctness*** | | | | |
| Task activity | L superior anterior temporal lobe | 0.25 | 0.008 (0.031) | 0.807 (0.852) |
| Task activity | R superior anterior temporal lobe | -1.26 | -0.042 (0.033) | 0.212 (0.636) |
| Task activity | B subgenual anterior cingulate cortex | -0.19 | -0.006 (0.031) | 0.852 (0.852) |
| Task-dependent connectivity | L superior anterior temporal lobe<br>B subgenual anterior cingulate cortex | 1.10 | 0.029 (0.027) | 0.275 (0.275) |
| Task-dependent connectivity | R superior anterior temporal lobe<br>B subgenual anterior cingulate cortex | -1.34 | -0.036 (0.027) | 0.186 (0.275) |
| ***Time-of-day*** | | | | |
| Task activity | L superior anterior temporal lobe | 0.16 | 0.004 (0.023) | 0.873 (0.873) |
| Task activity | R superior anterior temporal lobe | 1.57 | 0.038 (0.025) | 0.122 (0.366) |
| Task activity | B subgenual anterior cingulate cortex | -0.56 | -0.013 (0.023) | 0.577 (0.866) |
| Task-dependent connectivity | L superior anterior temporal lobe<br>B subgenual anterior cingulate cortex | 1.09 | 0.021 (0.020) | 0.281 (0.562) |
| Task-dependent connectivity | R superior anterior temporal lobe<br>B subgenual anterior cingulate cortex | -0.48 | -0.010 (0.020) | 0.636 (0.636) |
| ***Morningness-eveningness × distinctness*** | | | | |
| Task activity | L superior anterior temporal lobe | -1.02 | -0.008 (0.008) | 0.313 (0.470) |
| Task activity | R superior anterior temporal lobe | -0.16 | -0.001 (0.009) | 0.877 (0.877) |
| Task activity | B subgenual anterior cingulate cortex | -1.71 | -0.014 (0.008) | 0.093 (0.279) |
| Task-dependent connectivity | L superior anterior temporal lobe<br>B subgenual anterior cingulate cortex | 0.37 | 0.003 (0.007) | 0.712 (0.712) |
| Task-dependent connectivity | R superior anterior temporal lobe<br>B subgenual anterior cingulate cortex | 1.83 | 0.013 (0.007) | 0.073 (0.146) |

| | | | | |
|---|---|---|---|---|
| ***Morningness-eveningness × time-of-day*** | | | | |
| Task activity | L superior anterior temporal lobe | 0.02 | 0.000 (0.005) | 0.983 (0.983) |
| Task activity | R superior anterior temporal lobe | 0.03 | 0.000 (0.006) | 0.978 (0.983) |
| Task activity | B subgenual anterior cingulate cortex | -0.93 | -0.005 (0.005) | 0.354 (0.983) |
| Task-dependent connectivity | L superior anterior temporal lobe<br>B subgenual anterior cingulate cortex | -1.50 | -0.007 (0.004) | 0.139 (0.278) |
| Task-dependent connectivity | R superior anterior temporal lobe<br>B subgenual anterior cingulate cortex | 0.10 | 0.000 (0.005) | 0.921 (0.921) |
| ***Distinctness × time-of-day*** | | | | |
| ***Task activity*** | ***L superior anterior temporal lobe*** | ***-2.04*** | ***-0.017 (0.009)*** | ***0.046 (0.138)*** |
| Task activity | R superior anterior temporal lobe | -1.60 | -0.015 (0.009) | 0.115 (0.173) |
| Task activity | B subgenual anterior cingulate cortex | 0.19 | 0.002 (0.009) | 0.851 (0.851) |
| Task-dependent connectivity | L superior anterior temporal lobe<br>B subgenual anterior cingulate cortex | 1.88 | 0.014 (0.007) | 0.064 (0.128) |
| Task-dependent connectivity | R superior anterior temporal lobe<br>B subgenual anterior cingulate cortex | -0.18 | -0.001 (0.008) | 0.855 (0.855) |
| ***Morningness-eveningness × distinctness × time-of-day*** | | | | |
| Task activity | L superior anterior temporal lobe | 0.18 | 0.000 (0.002) | 0.862 (0.864) |
| ***Task activity*** | ***R superior anterior temporal lobe*** | ***-2.35*** | ***-0.004 (0.002)*** | ***0.022 (0.066)*** |
| Task activity | B subgenual anterior cingulate cortex | -0.17 | -0.000 (0.002) | 0.864 (0.864) |
| Task-dependent connectivity | L superior anterior temporal lobe<br>B subgenual anterior cingulate cortex | -0.44 | -0.000 (0.001) | 0.658 (0.658) |
| **Task-dependent connectivity** | **R superior anterior temporal lobe<br>B subgenual anterior cingulate cortex** | **2.72** | **0.004 (0.001)** | **0.008 (0.017)** |

Abbreviations: FDR, false discovery rate; L, left; R, right; B, bilateral; SE, standard error.

**Supplementary Table 5.** The main and interactions effects of morningness-eveningness, distinctness and time-of-day on the habituation of task-evoked activity and connectivity.

| Neural measure | Brain region | T-stat | β (SE) | $p_{uncorr.}$ ($p_{FDR}$) |
|---|---|---|---|---|
| ***Morningness-eveningness*** | | | | |
| Task activity | L superior anterior temporal lobe | 0.93 | 0.017 (0.018) | 0.357 (0.556) |
| Task activity | R superior anterior temporal lobe | 0.90 | 0.016 (0.018) | 0.373 (0.556) |
| Task activity | B subgenual anterior cingulate cortex | -0.59 | -0.010 (0.018) | 0.556 (0.556) |
| Task-dependent connectivity | L superior anterior temporal lobe<br>B subgenual anterior cingulate cortex | 1.20 | 0.023 (0.019) | 0.235 (0.570) |
| Task-dependent connectivity | R superior anterior temporal lobe<br>B subgenual anterior cingulate cortex | 0.57 | 0.010 (0.018) | 0.570 (0.570) |
| ***Distinctness*** | | | | |
| Task activity | L superior anterior temporal lobe | 0.23 | 0.005 (0.026) | 0.817 (0.817) |
| Task activity | R superior anterior temporal lobe | 0.31 | 0.008 (0.026) | 0.761 (0.817) |
| Task activity | B subgenual anterior cingulate cortex | 1.73 | 0.044 (0.026) | 0.089 (0.267) |
| Task-dependent connectivity | L superior anterior temporal lobe<br>B subgenual anterior cingulate cortex | -1.18 | -0.032 (0.028) | 0.224 (0.344) |
| Task-dependent connectivity | R superior anterior temporal lobe<br>B subgenual anterior cingulate cortex | -0.95 | -0.025 (0.026) | 0.344 (0.344) |
| ***Time-of-day*** | | | | |
| Task activity | L superior anterior temporal lobe | 0.17 | 0.003 (0.019) | 0.868 (0.950) |
| Task activity | R superior anterior temporal lobe | -0.06 | -0.001 (0.020) | 0.950 (0.950) |
| Task activity | B subgenual anterior cingulate cortex | -1.05 | -0.020 (0.019) | 0.298 (0.894) |
| Task-dependent connectivity | L superior anterior temporal lobe<br>B subgenual anterior cingulate cortex | 1.57 | 0.032 (0.020) | 0.123 (0.246) |
| Task-dependent connectivity | R superior anterior temporal lobe<br>B subgenual anterior cingulate cortex | 0.12 | 0.002 (0.020) | 0.906 (0.906) |
| ***Morningness-eveningness × distinctness*** | | | | |
| Task activity | L superior anterior temporal lobe | 0.46 | 0.003 (0.007) | 0.646 (0.865) |
| Task activity | R superior anterior temporal lobe | 0.28 | 0.002 (0.007) | 0.780 (0.865) |
| Task activity | B subgenual anterior cingulate cortex | 0.17 | 0.001 (0.007) | 0.865 (0.865) |
| Task-dependent connectivity | L superior anterior temporal lobe<br>B subgenual anterior cingulate cortex | 0.19 | 0.001 (0.007) | 0.852 (0.852) |
| Task-dependent connectivity | R superior anterior temporal lobe<br>B subgenual anterior cingulate cortex | 0.24 | 0.002 (0.007) | 0.809 (0.852) |
| ***Morningness-eveningness × time-of-day*** | | | | |
| Task activity | L superior anterior temporal lobe | 0.09 | 0.000 (0.004) | 0.928 (0.928) |

| | | | | |
|---|---|---|---|---|
| Task activity | R superior anterior temporal lobe | -0.88 | -0.004 (0.004) | 0.383 (0.651) |
| Task activity | B subgenual anterior cingulate cortex | 0.79 | 0.003 (0.004) | 0.434 (0.651) |
| Task-dependent connectivity | L superior anterior temporal lobe<br>B subgenual anterior cingulate cortex | 0.17 | 0.001 (0.005) | 0.864 (0.921) |
| Task-dependent connectivity | R superior anterior temporal lobe<br>B subgenual anterior cingulate cortex | 0.10 | 0.000 (0.004) | 0.921 (0.921) |
| ***Distinctness × time-of-day*** | | | | |
| Task activity | L superior anterior temporal lobe | -0.14 | -0.001 (0.007) | 0.887 (0.887) |
| Task activity | R superior anterior temporal lobe | 0.53 | 0.004 (0.007) | 0.597 (0.887) |
| Task activity | B subgenual anterior cingulate cortex | 0.68 | 0.005 (0.007) | 0.499 (0.887) |
| Task-dependent connectivity | L superior anterior temporal lobe<br>B subgenual anterior cingulate cortex | 0.03 | 0.000 (0.008) | 0.978 (0.978) |
| Task-dependent connectivity | R superior anterior temporal lobe<br>B subgenual anterior cingulate cortex | -0.14 | -0.001 (0.007) | 0.891 (0.978) |
| ***Morningness-eveningness × distinctness × time-of-day*** | | | | |
| Task activity | L superior anterior temporal lobe | -0.85 | -0.001 (0.001) | 0.398 (0.922) |
| Task activity | R superior anterior temporal lobe | -0.46 | -0.000 (0.001) | 0.649 (0.922) |
| Task activity | B subgenual anterior cingulate cortex | 0.10 | 0.000 (0.001) | 0.922 (0.922) |
| Task-dependent connectivity | L superior anterior temporal lobe<br>B subgenual anterior cingulate cortex | -0.09 | -0.000 (0.001) | 0.932 (0.932) |
| Task-dependent connectivity | R superior anterior temporal lobe<br>B subgenual anterior cingulate cortex | 0.35 | 0.000 (0.001) | 0.726 (0.932) |

Abbreviations: FDR, false discovery rate; L, left; R, right; B, bilateral; SE, standard error.

**Supplementary Table 6.** The main and interactions effects of morningness-eveningness, distinctness and time-of-day on the behavioural metrics associated with the autobiographical guilt memory recollection task. The results surviving the multiple comparison correction ($p_{FDR} < 0.05$) are shown in blue and bold, while nominally significant findings ($p_{uncorr.} < 0.05$) are additionally presented in italics.

| Variable | Morningness-eveningness | | Distinctness | | Time-of-day | | Morningness-eveningness × distinctness | | Morningness-eveningness × time-of-day | | Distinctness × time-of-day | | Morningness-eveningness × distinctness × time-of-day | |
|---|---|---|---|---|---|---|---|---|---|---|---|---|---|---|
| | T-stat | $p_{uncorr}$ ($p_{FDR}$) | T-stat | $p_{uncorr}$ ($p_{FDR}$) | T-stat | $p_{uncorr}$ ($p_{FDR}$) | T-stat | $p_{uncorr}$ ($p_{FDR}$) | T-stat | $p_{uncorr}$ ($p_{FDR}$) | T-stat | $p_{uncorr}$ ($p_{FDR}$) | T-stat | $p_{uncorr}$ ($p_{FDR}$) |
| ***Guilt memories*** | | | | | | | | | | | | | | |
| Strength of guilt feelings | 0.29 | 0.770 (0.974) | ***2.70*** | ***0.009 (0.077)*** | – | – | 1.93 | 0.058 (0.465) | – | – | – | – | – | – |
| Perceived social code violation | 0.72 | 0.477 (0.974) | 0.28 | 0.783 (0.783) | – | – | -0.09 | 0.930 (0.930) | – | – | – | – | – | – |
| Vividness outside the scanner | 0.54 | 0.599 (0.974) | 0.67 | 0.506 (0.735) | – | – | -1.08 | 0.277 (0.465) | – | – | – | – | – | – |
| Vividness inside the scanner | -0.03 | 0.974 (0.974) | ***2.10*** | ***0.039 (0.089)*** | 1.01 | 0.313 (0.470) | -0.50 | 0.625 (0.718) | -0.49 | 0.626 (0.920) | 1.25 | 0.220 (0.660) | 0.78 | 0.449 (0.607) |
| Approach-avoidance motivation | 0.11 | 0.910 (0.974) | -0.42 | 0.685 (0.771) | -1.02 | 0.313 (0.470) | 1.18 | 0.250 (0.465) | 0.79 | 0.435 (0.870) | -0.89 | 0.382 (0.764) | ***2.00*** | ***0.049 (0.293)*** |
| Salience | -1.41 | 0.157 (0.716) | ***2.11*** | ***0.039 (0.089)*** | 1.18 | 0.240 (0.470) | -1.03 | 0.306 (0.465) | -0.25 | 0.796 (0.920) | 1.56 | 0.126 (0.660) | 0.67 | 0.506 (0.607) |
| Accuracy for content-related questions | 0.44 | 0.662 (0.974) | 0.56 | 0.572 (0.735) | 1.27 | 0.205 (0.470) | -1.01 | 0.310 (0.465) | -0.10 | 0.920 (0.920) | -0.00 | 0.999 (0.999) | -0.89 | 0.375 (0.607) |
| Mean RT for content-related question | -0.37 | 0.707 (0.974) | ***2.35*** | ***0.022 (0.089)*** | -0.61 | 0.539 (0.647) | 1.04 | 0.304 (0.465) | 1.07 | 0.281 (0.843) | -0.14 | 0.886 (0.999) | -0.32 | 0.747 (0.747) |
| Successful recalls inside the scanner | 1.42 | 0.159 (0.716) | 0.73 | 0.466 (0.735) | 0.30 | 0.772 (0.772) | 0.48 | 0.638 (0.718) | -1.61 | 0.114 (0.684) | 0.59 | 0.557 (0.836) | 1.26 | 0.211 (0.607) |
| ***Neutral memories*** | | | | | | | | | | | | | | |
| Strength of guilt feelings | 0.54 | 0.599 (0.933) | -1.41 | 0.170 (0.493) | – | – | -0.25 | 0.795 (0.933) | – | – | – | – | – | – |
| Perceived social code violation | -0.04 | 0.964 (0.933) | -1.71 | 0.094 (0.493) | – | – | -0.39 | 0.665 (0.933) | – | – | – | – | – | – |
| Emotional valence | 0.97 | 0.334 (0.933) | -0.58 | 0.566 (0.773) | – | – | -0.15 | 0.880 (0.933) | – | – | – | – | – | – |
| Vividness outside the scanner | 0.39 | 0.706 (0.933) | -1.15 | 0.259 (0.518) | – | – | 0.08 | 0.933 (0.933) | – | – | – | – | – | – |
| Vividness inside the scanner | -0.13 | 0.897 (0.933) | -0.53 | 0.607 (0.773) | 0.79 | 0.434 (0.651) | 0.10 | 0.915 (0.933) | 0.53 | 0.597 (0.647) | -0.80 | 0.421 (0.819) | -0.09 | 0.922 (0.922) |

| Variable | Morningness-eveningness | | Distinctness | | Time-of-day | | Morningness-eveningness × distinctness | | Morningness-eveningness × time-of-day | | Distinctness × time-of-day | | Morningness-eveningness × distinctness × time-of-day | |
|---|---|---|---|---|---|---|---|---|---|---|---|---|---|---|
| | **T-stat** | **$p_{uncorr}$ ($p_{FDR}$)** | **T-stat** | **$p_{uncorr}$ ($p_{FDR}$)** | **T-stat** | **$p_{uncorr}$ ($p_{FDR}$)** | **T-stat** | **$p_{uncorr}$ ($p_{FDR}$)** | **T-stat** | **$p_{uncorr}$ ($p_{FDR}$)** | **T-stat** | **$p_{uncorr}$ ($p_{FDR}$)** | **T-stat** | **$p_{uncorr}$ ($p_{FDR}$)** |
| ***Neutral memories*** | | | | | | | | | | | | | | |
| Approach-avoidance motivation | 0.32 | 0.748 (0.933) | -0.50 | 0.618 (0.773) | 0.28 | 0.781 (0.781) | -0.38 | 0.708 (0.933) | 1.27 | 0.214 (0.351) | 0.54 | 0.587 (0.819) | 0.61 | 0.539 (0.922) |
| Salience | 0.81 | 0.430 (0.933) | -0.06 | 0.952 (0.952) | -1.45 | 0.148 (0.651) | 0.45 | 0.645 (0.933) | 1.24 | 0.222 (0.351) | -0.35 | 0.730 (0.819) | 0.40 | 0.693 (0.922) |
| Accuracy for content-related questions | 0.74 | 0.457 (0.933) | 1.32 | 0.197 (0.493) | 0.97 | 0.335 (0.651) | -0.62 | 0.534 (0.933) | -0.45 | 0.647 (0.647) | 1.24 | 0.220 (0.819) | -1.54 | 0.124 (0.744) |
| Mean RT for content-related question | -0.54 | 0.594 (0.933) | 0.27 | 0.779 (0.866) | 0.61 | 0.559 (0.671) | -0.33 | 0.731 (0.933) | 1.59 | 0.127 (0.351) | -0.23 | 0.819 (0.819) | 1.01 | 0.325 (0.922) |
| Successful recalls inside the scanner | -0.01 | 0.993 (0.933) | -1.35 | 0.176 (0.493) | 0.91 | 0.369 (0.651) | -1.79 | 0.078 (0.780) | 1.21 | 0.234 (0.351) | 1.09 | 0.277 (0.819) | -0.13 | 0.896 (0.922) |
| ***Guilt vs. neutral memories*** | | | | | | | | | | | | | | |
| Approach-avoidance motivation | -0.09 | 0.928 (0.928) | -0.09 | 0.925 (0.925) | -1.09 | 0.279 (0.578) | 1.28 | 0.206 (0.239) | -0.03 | 0.977 (0.977) | -1.12 | 0.267 (0.267) | 1.45 | 0.146 (0.292) |
| Salience | -1.82 | 0.076 (0.152) | 1.50 | 0.136 (0.272) | **2.35** | **0.025 (0.049)** | -1.19 | 0.239 (0.239) | -1.49 | 0.142 (0.284) | 1.43 | 0.158 (0.158) | 0.03 | 0.970 (0.970) |
| ***Distractor task*** | | | | | | | | | | | | | | |
| Accuracy | -0.07 | 0.949 (0.949) | 0.93 | 0.350 (0.350) | 0.24 | 0.809 (0.809) | 0.55 | 0.576 (0.576) | -0.19 | 0.847 (0.847) | -1.24 | 0.212 (0.424) | 0.18 | 0.844 (0.844) |
| Mean RT | -0.81 | 0.429 (0.858) | 1.05 | 0.310 (0.350) | 0.90 | 0.373 (0.746) | -0.96 | 0.339 (0.576) | ***2.15*** | ***0.035 (0.07)*** | 0.51 | 0.613 (0.613) | 0.23 | 0.819 (0.844) |

Abbreviations: FDR, false discovery rate; RT, reaction time.

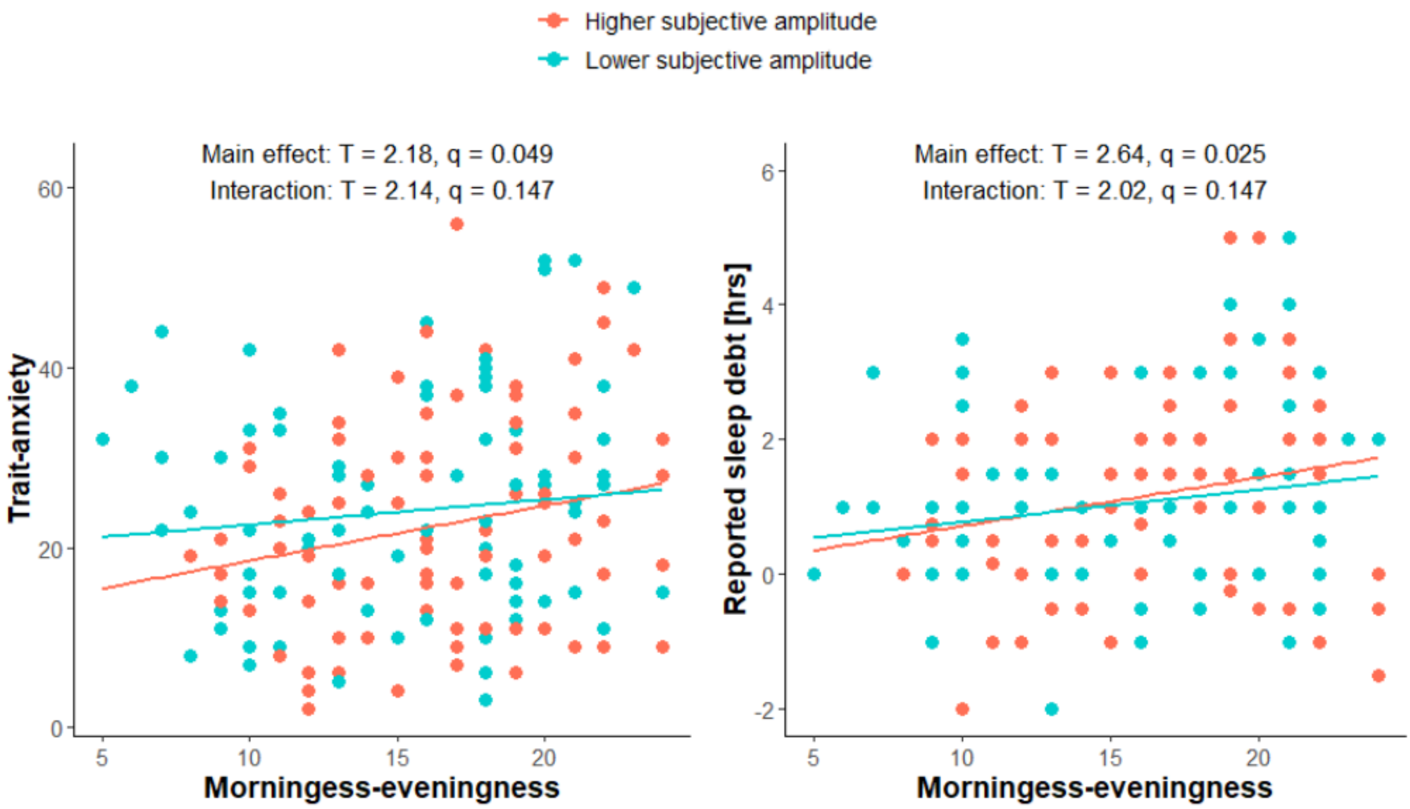


**Supplementary Figure S1.** Moderation effects of subjective diurnal rhythm amplitude (distinctness) on the associations of the morningness-eveningness scores with trait anxiety (left panel) and reported sleep debt (right panel). The division of the sample into the groups with lower (cyan) and higher (salmon) subjective amplitude was based on the median score within the study population (i.e. 13, with 25 being the highest possible value). Negative sleep debt values indicate that the individuals would typically sleep longer than the perceived minimum required to wake up feeling refreshed. Abbreviations: q, false discovery rate-corrected p value.

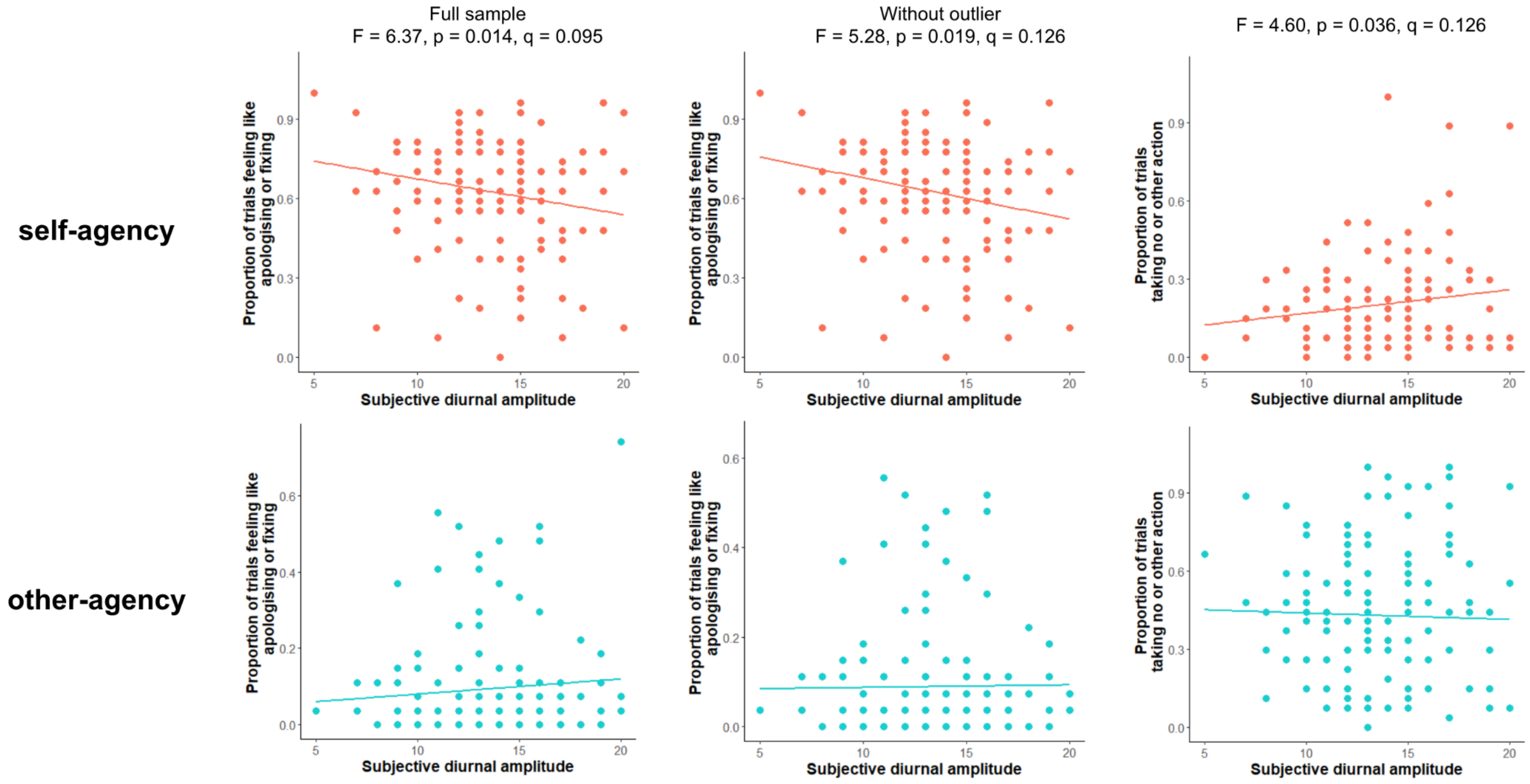


**Supplementary Figure S2.** Associations of subjective diurnal amplitude (distinctness) with action tendencies from the Moral Sentiment and Action Tendencies (MSAT) task. Top and bottom panels represent the data for self- and other-agency conditions, respectively. In the full sample, we observed a nominally significant interaction between distinctness and agency (first column), driven predominantly by the decreased proportion of self-agency trials where individuals with higher subjective diurnal amplitude felt like apologising or fixing what they had done. The second column represents the recalculated results, this time excluding the outlier visible in the original other-agency condition. The third column shows the nominally significant interaction between distinctness and agency with regards to choosing no or other action. Abbreviations: q, false discovery rate-corrected p value.

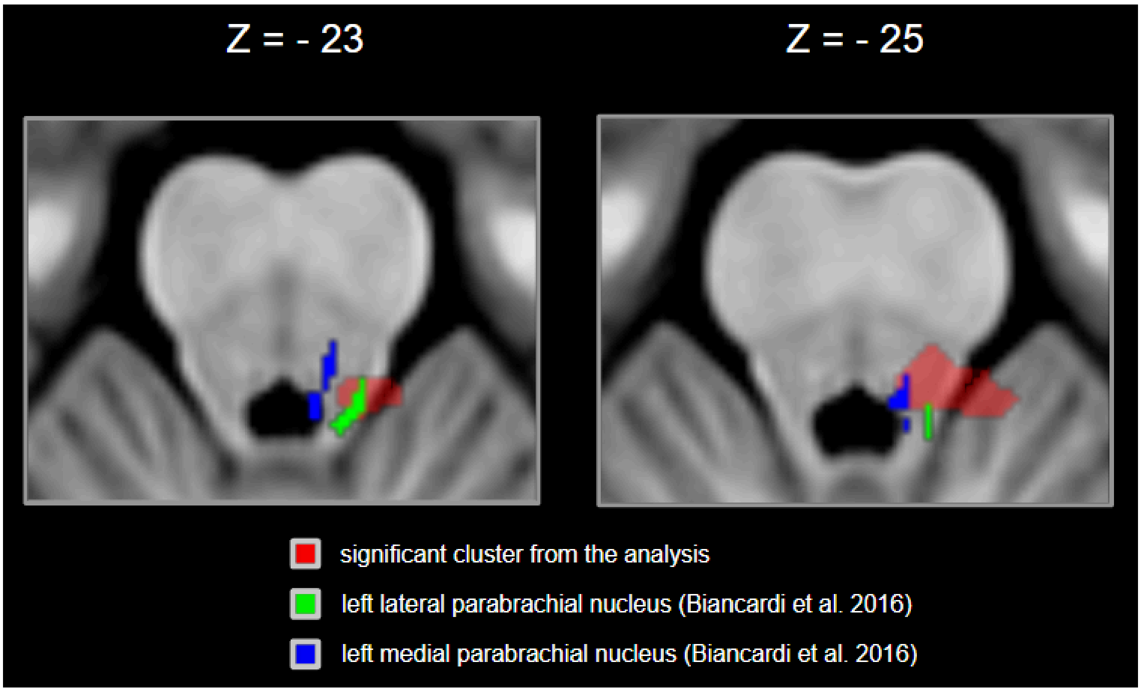


**Supplementary Figure S3.** Spatial overlap between the left medial and lateral parabrachial nuclei, as delineated with a 7T brainstem atlas (Biancardi et al., 2016), and the cluster with stronger habituation of the right superior anterior temporal lobe task-related connectivity in individuals with higher distinctness.
